# Not just BLAST nt: WGS database joins the party

**DOI:** 10.1101/653592

**Authors:** Jose Manuel Martí, Carlos P. Garay

**Affiliations:** Institute for Integrative Systems Biology(I^2^SysBio), Valencia, Spain; Canfranc Underground Laboratory (LSC), Huesca, Spain

**Keywords:** metagenomics, taxonomic classification, draft genomes, whole genome, NCBI, GenBank, WGS, sensitivity

## Abstract

Since its introduction in 1990 and with over 50k citations, the NCBI BLAST family has been an essential tool of *in silico* molecular biology. The BLAST `nt` database, based on the traditional divisions of GenBank, has been the default and most comprehensive database for nucleotide BLAST searches and for taxonomic classification software in metagenomics. Here we argue that this is no longer the case. Currently, the NCBI WGS database contains one billion reads (almost five times more than GenBank), and with 4.4 trillion nucleotides, WGS has about 14 times more nucleotides than GenBank. This ratio is growing with time. We advocate a change in the database paradigm in taxonomic classification by systematically combining the `nt` and WGS databases in order to boost taxonomic classifiers sensitivity. We present here a case in which, by adding WGS data, we obtained over five times more classified reads and with a higher confidence score. To facilitate the adoption of this approach, we provide the `draftGenomes` script.

**Author summary:** Culture-independent methods are revolutionizing biology. The NIH/NCBI Basic Local Alignment Search Tool (BLAST) is one of the most widely used methods in computational biology. The BLAST `nt` database has become a de facto standard for taxonomic classifiers in metagenomics. We believe that it is time for a change in the database paradigm for such a classification. We advocate the systematic combination of the BLAST `nt` database with genomes of the massive NCBI Whole-Genome Shotgun (WGS) database. We make `draftGenomes` available, a script that eases the adoption of this approach. Current developments and technologies make it feasible now. Our recent results in several metagenomic projects indicate that this strategy boosts the sensitivity in taxonomic classifications.

## 1 Introduction

The NIH (National Institute of Health) created the NCBI (National Center for Biotechnology Information) in 1988 to advance information systems devoted to molecular biology. The GenBank nucleic acid sequence database was one of its first hosted projects (Benson et al. 2013). When the NCBI Taxonomy database project was launched in 1991, GenBank was moved under its umbrella (Federhen 2011). Nowadays, NCBI provides computational resources and data retrieval systems for the study of several other sets of biological data arranged in various additional databases (NCBI Resource Coordinators 2013). The NCBI BLAST (Basic Local Alignment Search Tool) software (Altschul et al. 1990) uses a standard set of BLAST databases for protein, nucleotide, and translated BLAST searches. NCBI frequently releases pre-formatted versions of these databases as compressed files, which any user can download from the NCBI BLAST FTP site. The NCBI BLAST `nt` database is one of those databases and is actually the default for nucleotide BLAST searches (NCBI Resource Coordinators 2013).

Common misunderstandings about the BLAST `nt` database include the belief that it is a strictly non-redundant nucleotide sequence database, and that all sequences from the NCBI nucleotide databases are included. Currently (as of May 2019), it contains “*partially non-redundant nucleotide sequences from all traditional divisions of GenBank, EMBL, and DDBJ; excluding GSS, STS, PAT, EST, HTG, and WGS*” (NCBI 2017). Therefore, while `nr` is strictly non-redundant, `nt` is partially non-redundant (NCBI 2017).

Concretely, the BLAST `nt` database includes the NCBI nucleotide sequences databases listed in Table 1 but does not include the NCBI nucleotide sequences databases indexed in Table 2, which also details how to obtain those resources downloadable from NCBI.

**Table 1.** NCBI nucleotide sequences databases included in BLAST `nt` database.

| NCBI database | Contents |
| --- | --- |
| TSA | Transcriptome shotgun data |
| ENV | Environmental samples |
| PHG | Phages |
| BCT | Bacteria |
| INV | Invertebrates |
| VRL | Viruses |
| MAM | Other mammals |
| PLN | Plants |
| SYN | Synthetic |
| VRT | Other vertebrates |
| UNA | Unannotated |
| PRI | Primates |
| ROD | Rodents |
| HTC | High-throughput cDNA |

**Table 2.** NCBI nucleotide sequences databases NOT included in BLAST `nt` database.

| NCBI DB | Contents | Additional files to download from NCBI FTP servers |
| --- | --- | --- |
| GSS | Genome survey sequences | gss.*tar.gz |
| STS | Sequence tagged sites | sts.gz |
| PAT | Patented sequences | patnt.gz |
| EST | Expressed sequence tags | est_human.gz,<br>est_mouse.gz,<br>est_others.gz |
| HTC | High-throughput cDNA | htgs.*tar.gz |
| WGS | Whole-genome shotgun data | Use draftGenomes |

Notice that the `nt` database includes genomes sequenced by a whole genome shotgun strategy but not the WGS database, a massive database from NCBI (Wheeler et al. 2007) that contains incomplete genomes sequenced by whole genome shotgun. These nucleotide sequences belong to hundreds of thousands of different sequencing projects which should be located and downloaded individually. Precisely, `draftGenomes` understands this project-basis approach and automates the collection of all the sequences in the NCBI WGS database belonging to a taxonomic subtree.

Strictly speaking, the WGS database is a division of the GenBank database that appeared in April 2002, 21 years after the first release of GenBank. Nevertheless, the name ‘GenBank’ is often used when referring to the traditional GenBank divisions, which do not include the WGS database. We will keep this tacit agreement throughout this manuscript and will consider GenBank and WGS as independent databases.

In August 2005, after only 3 years and four months in use, the WGS database surpassed GenBank in number of nucleotides. Years later, in June 2014, the release 202 of the WGS database exceeded GenBank also in the number of sequences included. In recent years, the number of bases and sequences in the WGS database is roughly doubling every two years.

Figure 1 shows the evolution of both the number of bases and the number of sequences (in semilogarithmic scale) contained in the GenBank and WGS databases, from December 1982 to April 2019. Interestingly, the GenBank database presented an exponential growth in the early stages, very similar to the current increase rate of the WGS database. In recent years, the growth of the GenBank database has slowed down significantly. The number of GenBank bases increased threefold in the last decade, far from the 48-fold increase that took place from 1998 to 2008.

**Figure 1.**
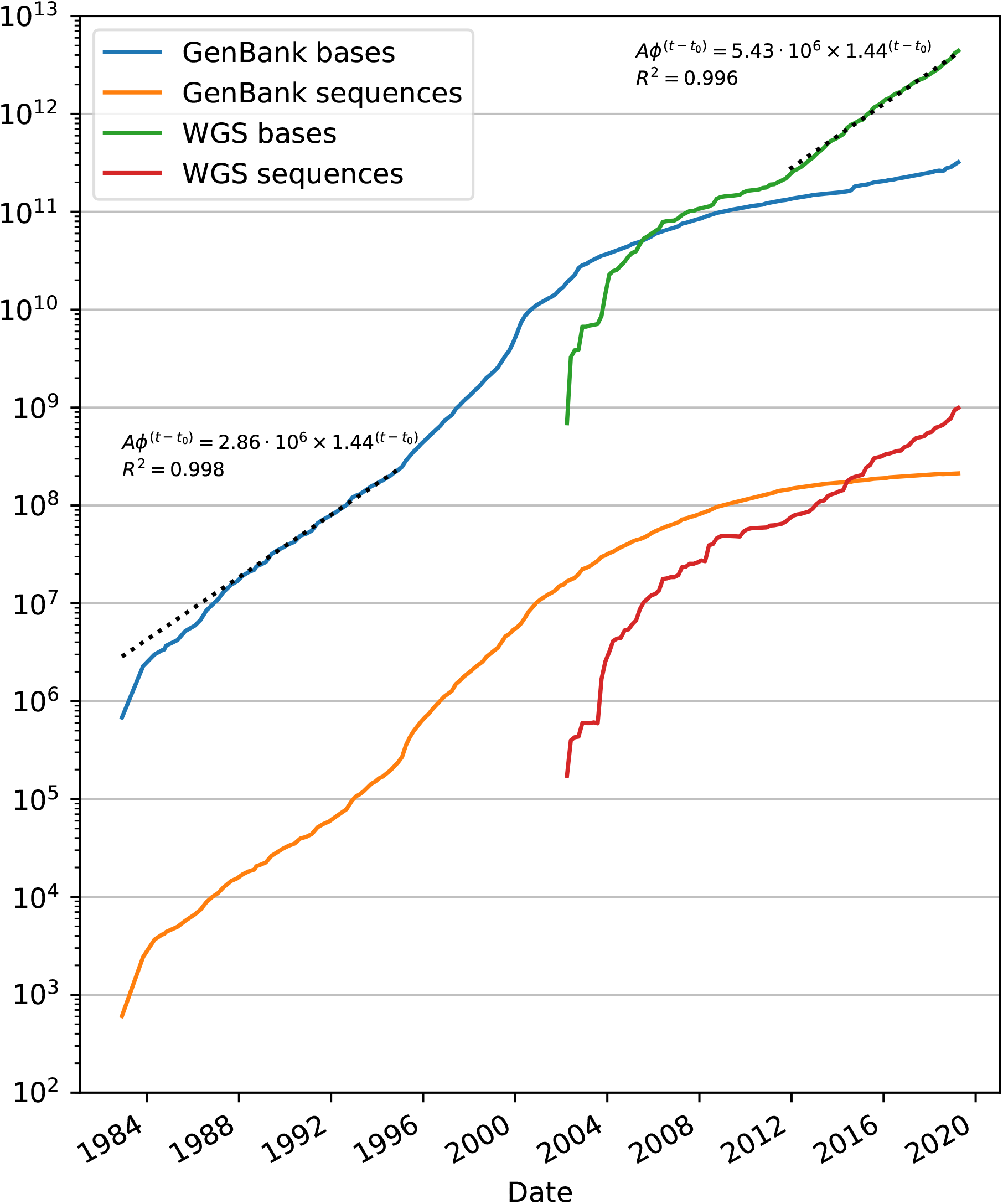
Temporal evolution of the NCBI GenBank and WGS databases from December 1982 to April 2019. The y-axis indicates the number of bases and sequences (in logarithmic scale) and the x-axis the time in years (with December 1982 as the initial time *t*_0_). Additionally, the figure shows as dotted black lines the fits of exponential models to the number of bases. Interestingly, from the beginning to December 1994, GenBank shows an exponential growth rate in the number of bases very similar to that of WGS from December 2011 to the last release (April 2019).

## 2 Design and implementation

### 2.1 Overview

The `draftGenomes` software greatly simplifies the otherwise arduous task of collecting all the NCBI WGS sequences related to a NCBI taxonomic identifier (TaxID) at any taxonomic level. This script downloads the appropriate sequence files from NCBI WGS projects and parses them to generate a single coherent fasta file by parsing the sequence headers and updating them if needed.

### 2.2 Details

Depending on the chosen TaxID, the download and analysis of the NCBI WGS projects can take a long time (and require a lot of disk space). To deal with such a common situation, the script shows progress indicators and automatically recovers from several errors. Furthermore, `draftGenomes` has been fitted with a `resume mode` in case of any fatal interruption of the process.

We describe below further modes of operation:

- The `reverse mode` enables another instance of the script to manage the download of sequences in reverse order without interfering with the first one, which is also parsing the sequences to generate the resulting fasta file.
- The `force mode` ignores previous downloads and recreates the final FASTA file in spite of previous runs of the script for the same TaxID.
- The `download mode` is used to download without parsing the WGS project files.
- The `verbose mode` substitutes the progress indicator with details about every project parsed.

The `draftGenomes` code has been tested successfully in ~TB downloads from the NCBI servers with several forced and unforced interruptions.

### 2.3 Installing draftGenomes

`draftGenomes` is a compact script that can be installed just cloning the GitHub repository, downloading the script, or even pasting the source code using any text editor.

### 2.4 Running draftGenomes

`draftGenomes` just requires a Python 3 interpreter. No other packages beyond the Python Standard Library ones are needed. Running. /draftGenomes.py --help will print all the possibilities and details of the `draftGenomes` software. The output files have the format: `WGS4taxid{include}-{exclude}.fa` where `{include}` is the TaxID of the root of the taxonomic subtree of interest while `{exclude}` (optional) is the TaxID of the root of the excluded taxa in that subtree. Both TaxIDs are options of the script (a run with no TaxID related arguments will test the script).

### 2.5 Using draftGenomes with Centrifuge and Recentrifuge

Centrifuge’s (Kim et al. 2016) `nt`+WGS databases are the result of pre-processing the NCBI BLAST `nt` database plus NCBI WGS sequences collected by `draftGenomes` so that they can be used by Centrifuge, thus allowing rapid and very sensitive classification within a huge range of organisms. Recentrifuge (Martí 2019) is able to post-process the results of Centrifuge (and several other classification engines) and provide an interactive, score-oriented visualization for the final outcome.

## 3 Results

In Martí et al. (2019), we prepared (December 2017) an ad hoc database for the Centrifuge classifier (Kim et al. 2016) including all the sequences from the `nt` database and some others from the NCBI WGS database obtained with `draftGenomes`. Those extra sequences belonged to the genus *Olea* and the kingdom *Fungi*. We used the resulting Centrifuge database to analyze several samples from a transcriptomic study of the evolution of Verticillium wilt of olive (Jimenez-Ruiz et al. 2017).

Figure 2 shows a comparison of results with the `nt` versus `nt`+WGS databases for the sample corresponding to olive tree roots 15 days after *Verticilium dahliae* inoculation. We see in the figure that, with the `nt`+WGS database, the number of sequences classified passing the score filter was more than five times higher. The use of the `nt` database led to a final classification rate of 6.4% of the sequenced reads, whereas this rate increased to 34.7% when the `nt` database was used together with the chosen sequences from the WGS database. We selected a high value (60 bp) as the threshold level for the score filter as recommended for RNA-sequencing projects (Kim et al. 2016).

**Figure 2.**
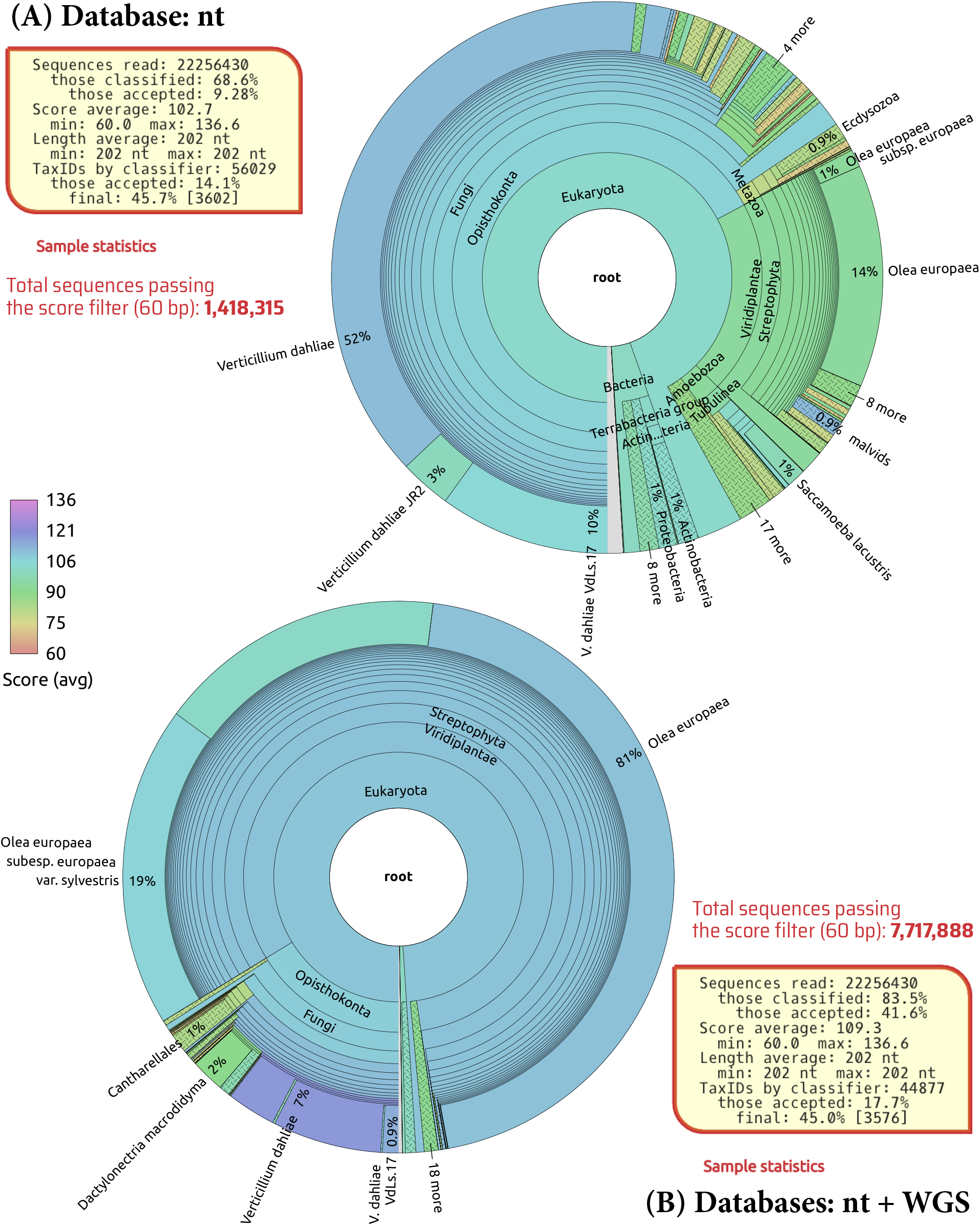
Comparison of results with the nt database versus sequences of WGS on top of the nt database. Recentrifuge (Martí 2019) plots of SMS classified reads by Centrifuge, (**A**) using the `nt` database and (**B**) using the `nt` database plus all the sequences of the genus *Olea* and the kingdom *Fungi* present in the NCBI WGS database (Martí et al. 2019). The sample corresponds to Olive tree roots 15 days after *Verticilium dahliae* inoculation (Jimenez-Ruiz et al. 2017). The databases were retrieved and prepared in December 2017.

Nevertheless, the increase in the classification rate was not at the expenses of a decrease in the confidence score. On the contrary, the average confidence score grew from 102.7 to 109.3. Interestingly, as Figure 2 also shows, the number of different NCBI taxonomic identifiers (`TaxIDs` field of the sample statistics) that were hit by the classifier also decreased with the addition of WGS data, especially those associated with low-quality scores. Therefore, the addition of draft genomes related to the host (the olive tree) not only markedly increased the number of previously unclassified reads that were finally classified as belonging to the host, but also reduced the number of previously misclassified reads, which ultimately were assigned to the olive tree as well. Specifically, the number of reads classified into the *Olea europaea* TaxID or below had a 31-fold jump, and the score increased from 93.9 to 110.1, thus indicating that not only one but both sequences of the paired-end reads were partially matching.

Although the main effect of adding WGS data affected the detection of the host, the addition of draft fungal genomes provided some improvements too. For example, *Fusarium oxysporum*, which is considered the pathogenic agent of Fusarium wilt of chickpea (García-Pedrajas et al. 1999), had a 27.6% rise in the number of reads assigned and a 24.4% enhancement in the average confidence, to a relatively very high score of 120.5.

## 4 Discussion

Although the idea of using draft genomes to improve the classification sensitivity is not new (Ames et al. 2015), we find in our studies (Martí et al. 2019; Bernabeu-Gimeno et al. 2018) that the WGS database is mature and rich enough to support a systematic use of its projects to supplement the `nt` database, thus minimizing the time gap between partial sequencing of new species and their identification in worldwide samples.

Finally, new algorithms for removing contaminants and low-complexity sequences from databases of draft genomes (Lu and Salzberg 2018) strengthen our proposal. Our suggestion implies large customized databases reduced to the size of ~100 GB using compression algorithms, currently feasible in fat computing nodes. The emergence of SCM (Storage Class Memory) will shortly democratize high-performance data analytics (Weiland et al. 2018), a new model that will allow huge metagenomic databases to be recreated with fresh WGS data in an ordinary computer, just before their use.

### Availability

The data and source code are anonymously and freely available on GitHub at https://github.com/khyox/draftGenomes. The `draftGenomes` code is licensed under the GNU Affero General Public License Version 3 (https://www.gnu.org/licenses/agpl.html). The *readme* file of the GitHub’s repository is the most extensive and updated source of documentation for `draftGenomes`.

## Acknowledgments

We would like to thank the SOM group (IFIC, Spain) for some high performance computing and storage resources that we used in our tests of `draftGenomes`.

## Author Contributions

**Conceptualization:** JMM.

**Data curation**: JMM.

**Formal analysis**: JMM.

**Investigation**: JMM.

**Methodology**: JMM.

**Project administration**: JMM.

**Resources**:

**Software**: JMM.

**Supervision**: CPG.

**Validation**: JMM and CPG.

**Visualization**: JMM.

**Writing — original draft**: JMM.

**Writing — review & editing**: JMM and CPG.

## Supplemental Material

## Appendix A

### Abbreviations

BLAST: Basic Local Alignment Search Tool
DDBJ: DNA Data Bank of Japan
EMBL: European Molecular Biology Laboratory
EST: Expressed Sequence Tags NCBI database
GSS: Genome Survey Sequences NCBI database
HTG: High-throughput unfinished genome sequences NCBI database
NCBI: National Center for Biotechnology Information
NIH: National Institute of Health
nt: Partially non-redundant nucleotide sequences NCBI database
PAT: Patented sequences NCBI database
SCM: Storage Class Memory
STS: Sequence Tagged Sites NCBI database
WGS: Whole-Genome Shotgun NCBI database

## Appendix B: Step by step instructions on the use of draftGenomes in the generation of a custom nt+WGS database for Centrifuge

Currently, we need high-performance computing resources available to be able to generate the Centrifuge `nt`+WGS kind of databases. Typically, a *fat-node* will do the job (we used, at least, a 32 cores machine with half a tebibyte memory and a fast scratch storage system). These are the detailed instructions, which assume that you have Centrifuge (Kim et al. 2016) already installed and available in your system:

1. Clone the `draftGenomes` GitHub repository, for example, under the user’s home directory.

~~~
cd ~
git clone https://github.com/khyox/draftGenomes.git
~~~
2. The following action is to download the NCBI `nt` database and then to unzip it (both operations take some time as it has several tens of GiB):

~~~
mkdir nt_WGS
cd nt_WGS
wget ftp://ftp.ncbi.nih.gov/blast/db/FASTA/nt.gz
gunzip nt.gz
mv -v nt nt.fa
~~~
3. Proceed analogously with the `taxdump` databases (this is the shorter step):

~~~
mkdir taxonomy
cd taxonomy
wget ftp://ftp.ncbi.nlm.nih.gov/pub/taxonomy/taxdump.tar.gz
tar xvzf taxdump.tar.gz
cd . .
~~~
4. Use `draftGenomes` to automatically retrieve and consolidate the sequences of interest from the NCBI WGS database. For example, to obtain all the sequences associated with the genus *Olea* (NCBI TaxID 4145) and below, issue the following commands:

~~~
~/draftGenomes/draftGenomes -t 4145
mv -v WGS4taxid4145.fa Olea.fa
~~~
5. Repeat the previous step as needed if you would like `draftGenomes` to obtain different sets of sequences belonging to subtrees of unrelated TaxIDs (not intersecting, towards more specific levels of the taxonomic tree).
6. Now, to properly generate the accession to TaxID mapping file (that will be ~10 GiB), the following commands are needed:

~~~
wget ”ftp://ftp.ncbi.nih.gov/pub/taxonomy/accession2taxid/nucl_*.accession2taxid.gz”
wget ftp://ftp.ncbi.nih.gov/pub/taxonomy/accession2taxid/pdb.accession2taxid.gz
gunzip -c *.accession2taxid.gz | awk -v OFS=’ 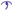 ’print $2, $3’ >> acc2tax.map
~~~
7. This is an optional step that will mask low-complexity sequences by using DustMasker (Morgulis et al. 2006), a NCBI BLAST command-line application that should be installed in the system (with the rest of the NCBI tools or alone from here). The code `dustmasker` should be executed with the DUST level (score threshold for subwindows) set to 20, which is the default. Finally, all the masked nucleotides from the DustMasker output will be remasked as `N` using sed:

~~~
mv nt.fa nt_unmasked.fa
dustmasker-infmt fasta -in nt_unmasked.fa -level 20 -outfmt fasta | sed ’/^>/! s/[^AGCT]/N/g’ > nt.fa
~~~
8. This is another optional step, and consists in repeating the last step but for any of the databases generated by `draftGenomes`. Just substitute the `nt` name for the applicable one, such as `Olea` for our example above.
9. Last but not least, the Centrifuge command that will generate the Centrifuge `nt`+WGS database (this is the part that actually benefits from high performance computing) must be issued:

~~~
centrifuge-build --ftabchars=14 -p 32 --bmax 1342177280 --conversion-table acc2tax.map
--taxonomy-tree taxonomy/nodes.dmp --name-table taxonomy/names.dmp nt.fa,Olea.fa nt_Olea
~~~

The last process finishes with a line like the following:

~~~
Total time for call to driver() for forward index: HH:MM:SS
~~~

That is the time of the last phase. In our case (using 32 cores) it took more than 20 hours. As this is not a short time, if `centrifuge-build` is launched not using a batch system but in an interactive session, we strongly recommend using any mechanism to protect the process from unintentional interruptions, for instance by `nohup`:

~~~
nohup centrifuge-build --ftabchars=14 -p 32 --bmax 1342177280 --conversion-table acc2tax.map
--taxonomy-tree taxonomy/nodes.dmp --name-table taxonomy/names.dmp nt.fa,Olea.fa nt_Olea &
~~~

In this case, use **tail -f nohup.out** to safely follow the progress of the `nt`+WGS database build.

